# The BSGatlas: An enhanced annotation of genes and transcripts for the *Bacillus subtilis* genome with improved information access

**DOI:** 10.1101/807263

**Authors:** Adrian Sven Geissler, Christian Anthon, Enrique González-Tortuero, Line Dahl Poulsen, Thomas Beuchert Kallehauge, Stefan Ernst Seemann, Jeppe Vinther, Jan Gorodkin

## Abstract

The genome of *Bacillus subtilis* continues to provide exiting genomic insights. However, the growing collective genomic knowledge about this micro-organism is spread across multiple annotation resources. Thus, the full annotation is not directly accessible neither for specific genes nor for large-scale high-throughput analyses. Furthermore, access to annotation of non-coding RNA genes (ncRNAs) and polycistronic mRNAs is difficult. To address these challenges we introduce the *Bacillus subtilis* genome atlas, BSGatlas, in which we integrate and unify multiple existing annotation resources. Our integration provides twice as many ncRNAs than the individual resources, improves the positional annotation for 70% of the combined ncRNAs, and makes it possible to infer specific ncRNA types. Moreover, we unify known transcription start sites, termination, and transcriptional units (TUs) as a comprehensive transcript map. This transcript map implies 815 new TUs and 6, 164 untranslated regions (UTRs), which is a five-fold increase over existing resources. We furthermore, find 2, 309 operons covering the transcriptional annotation for 93% of all genes, corresponding to an improvement by 11%. The BSGatlas is available in multiple formats. A user can either download the entire annotation in the standardized GFF3 format, which is compatible with most bioinformatics tools for omics and high-throughput studies, or view the annotation in an online browser at http://rth.dk/resources/bsgatlas.

**Importance:** The *Bacillus subtilis* genome has been studied in numerous context and consequently multiple efforts have been made in providing a complete annotation. Unfortunately, a number of resources are no longer maintained, and (i) the collective annotation knowledge is dispersed over multiple resources, of which each has a different focus of what type of annotation information they provide. (ii) Thus, it is difficult to easily and at a large scale obtain information for a genomic region or genes of interest. (iii) Furthermore, all resources are essentially incomplete when it comes to annotating non-coding and structured RNA, and transcripts in general. Here, we address all three problems by first collecting existing annotations of genes and transcripts start and termination sites; afterwards resolving discrepancies in annotations and combining them, which doubled the number of ncRNAs; inferring full transcripts and 2,309 operons from the combined knowledge of known transcript boundaries and meta-information; and critically providing it all in a standardized UCSC browser. That interface and its powerful set of functionalities allow users to access all the information in a single resource as well as enables them to include own data on top the full annotation.

---

*Bacillus* (Firmicutes, Bacilli) are Gram-positive soil microorganisms, which are used in biotech industrial applications, such as the industrial production of proteins, including enzymes^1^. *B. subtilis* is, besides *Escherichia coli*, the best studied bacterial specie. The application of modern high-throughput omics methods, such as RNA-seq^2, 3^, has provided important insight into *B. subtilis* gene regulation and demonstrated that bacterial transcription is more complex than previously expected^4, 5^. However, the interpretation of these analyses is highly dependent on the quality of annotation. Therefore, the *B. subtilis* genome annotation is subjected to ongoing, manual expert curation^6^, which requires substantial and non-trivial efforts^6, 7^. In spite of all this, the current annotation of *B. subtilis* remains highly protein-centric, such that the focus is on genomic elements related to messenger RNAs (mRNAs) and proteins^8^, whereas less attention is paid to other genomic elements, such as non-coding RNA genes and RNA structures (ncRNAs). The wide variety of ncRNA classes have diverse biological functions. For instance, the ribosomal RNA (rRNA) and transfer RNA (tRNA) are central to the protein translation process^9^, and the transfer messenger RNA (tmRNA) can rescue stalled ribosomes^10^. Moreover, *B. subtilis* has multiple ncRNAs and RNA structures that can regulate protein translation via interaction with mRNAs: Multiple mRNAs themselves contain cis-regulatory non-coding elements (e.g. riboswitches) that regulates the translation by limiting the access of the ribosome to the Shine-Dalgarno sequence^11^. In addition, small regulatory RNAs (sRNAs) can hybridize with mRNAs and thereby impact translational efficiency or mRNA stability^12^. A special case of sRNAs are ncRNAs that are expressed antisense to an mRNA, which may have large impact on the expression of the gene in the sense direction^13^.

Bacterial mRNAs differ from eukaryotic mRNA in the sense that they generally do not have splicing (although there are few occurrences of self-splicing intron structures^14, 15^), and that they can be poly-cistronic. Thus, *B. subtilis* mRNAs do not only have untranslated regions (UTRs) on their 5’ and 3’ ends, they also have internal UTRs^16^. Further adding to the complexity, bacterial operons can have multiple isoforms, due to the existence of alternative promoters and termination sites^4, 16^. The transcription of *B. subtilis* mRNAs are controlled by transcriptional factors that bind the DNA and promote or inhibit the transcription^16–18^. Yet, the termination of transcription depends on the presence of terminator hairpins or the binding of the *rho* protein to the nascent RNA^19^.

There exist multiple resources of *B. subtilis* genome annotation which are either individually redundant or contain mutually exclusive information (Table 1). This information range from high-quality curated information, experimental data to high-throughput based annotations. Hence, to fully exploit the available information of *B. subtilis* it is therefore necessary to fuse these into a single resource. These resources differ in their focus with some annotations attempting to narrow down the accurate genome coordinates of genes, operons, UTRs, and other genomic elements, while others focus on the annotation of biological functions, mutual interactions, and other meta information The RefSeq reference genome of *B. subtilis* strain 168^20, 21^ is the standard genome annotation relative to which other many experimental studies state their gene coordinates^5, 6, 22^. This reference annotation is also the basis for many other database and online analysis resources use either the identical or a very similar annotation^8, 23–26^ Besides *B. subtilis* specific or general genome annotation resources, ncRNA specific databases exist, which also relate to the RefSeq reference^27^. Due to the common dependency on the RefSeq reference, the number of resources that provide non-redundant information is comparably low. Thus, a focus on these resources, which we describe below, plays to the tasks of generating a single resources.

**Table 1.** Non-exhaustive, yet comprehensive overview over *B. subtilis* specific, general bacterial, or ncRNA focused annotation resources. Given is a short description for each resources, and what kind of annotations it provide. Resources noted with a check mark were considered for integration into the BSGatlas.

| Resource | General Description | Provided annotation |
| --- | --- | --- |
| RefSeq<sup>20</sup> ✓ | - Large collection of reference sequences incl. annotations<br>- Contains *B. subtilis* 168 standard annotation<br>- Most commonly used *B. subtilis* annotation resource | - Coding genes<br>- All rRNAs, tRNAs<br>- Only few other ncRNAs<br>- No mRNA transcripts |
| DBTBS<sup>23</sup> ✓ | - Collection of transcriptional binding factors<br>- Literature curated database | - Operons<br>- TSS / TTS |
| BsubCyc<sup>31</sup> ✓ | - *B. subtilis* specific database<br>- Encyclopedia of metabolism and pathways | - Coding genes<br>- All rRNAs, tRNAs<br>- Only few other ncRNAs<br>- Some transcripts with TSS and TTS |
| SubtiWiki<sup>8</sup> ✓ | - *B. subtilis* specific database<br>- Active community annotation effort<br>- Functional annotation driven<br>- Does not provide coordinates | - Coding genes<br>- All rRNAs, tRNAs<br>- Many additional ncRNAs<br>- Transcriptional Units |
| Nicolas *et al.*<sup>5</sup> ✓ | - Tilling-array study in *B. subtilis*<br>- Transcriptome in over >100 conditions<br>- Large number of predictions<br>- Includes high-trust annotation by original author curation<br>- Technical resolution limitation | - Many predicted ncRNAs<br>- Many predicted TSS/TTS<br>- Explicitly annotates UTRs |
| Dar *et al.*<sup>22</sup> ✓ | - term-seq study in *B. subtilis* | - Riboswitches |
| Rfam<sup>33</sup> ✓ | - Database of RNA structure families<br>- We scanned *B. subtilis* genome with most recent Rfam version<sup>34</sup><br>- Database of RNA structure families | - All rRNAs, tRNAs<br>- Riboswitches<br>- sRNAs and other structured RNAs |
| Ensembl<sup>24</sup> | - Large collection of reference sequences incl. annotations<br>- Fully contained in RefSeq annotation | - Coding genes<br>- All rRNAs, tRNAs<br>- Only few other ncRNAs<br>- No mRNA transcripts |
| PATRIC<sup>25</sup> | - Integrative database of bacterial genomes<br>- Analysis tools to support biomedical research<br>- Uses different genome sequence then RefSeq<br>- But nearly identical on annotated gene set | - Coding genes<br>- All rRNAs, tRNAs<br>- Only few other ncRNAs<br>- No mRNA transcripts |
| IMG<sup>26</sup> | - Integrative database of Archaea, Bacteria, and Eukarya<br>- Tools for comparative genome analysis<br>- Uses RefSeq as reference | - Coding genes<br>- All rRNAs, tRNAs<br>- Only few other ncRNAs<br>- No mRNA transcripts |
| BSRD<sup>27</sup> | - Single ncRNA type specific database<br>- Already included in other resources | - sRNA |
| SRPDB<sup>28</sup> | - Single ncRNA type specific database<br>- Already included in other resources | - SRP |
| tmRDB<sup>28</sup> | - Single ncRNA type specific database<br>- Already included in other resources | - tmRNA |
| tmRNA<sup>29</sup> | - Single ncRNA type specific database<br>- Already included in other resources | - tmRNA |
| tRNAdb<sup>30</sup> | - Single ncRNA type specific database<br>- Already included in other resources | - tRNA |

The database of transcriptional regulation in Bacillus subtilis (DBTBS^23^) contains annotation of transcript factor binding sites, transcription start sites (TSSs), transcription termination sites (TTSs), and operons that have been extracted from hundreds of literature references. The annotation of DBTBS is partially contained in the curated databases BsubCyc, another high-quality resource, although BsubCyc focuses more on gene annotation, including an impressive amount of meta-information.^31^. Additionally, BsubCyc provides gene functions, also in the form of Gene Ontology (GO) terms^32^, and transcriptional regulatory relationships, and a fine-grained, stoichiometric annotation of enzymatic reactions and molecular interactions, which are relevant for metabolomic studies. However, both DBTBS and BsubCyc are discontinued projects. In contrast, SubtiWiki is a still active community-driven project focused on *B. subtilis* annotation^8^. Currently, SubtiWiki is the most commonly used database for obtaining gene information in the *B. subtilis* community^6^.

SubtiWiki provides abundant meta-information, including an organism-specific gene categorization, lists of transcriptional regulations and interactions, and description of transcriptional units (TUs). In fact, SubtiWiki already included some parts of BsubCyc, but not the curated list of TSSs, TTSs, and functional annotation via GO terms. Additionally, SubtiWiki provides a custom gene categorization, which has been used in the functional analysis of *B. subtilis* gene expression in RNA-seq data^3^. A recent expert gene annotation refinement from February 2018 has been included^6^, and the RefSeq reference annotation fully includes this refinement. Overall, SubtiWiki focuses on meta-information instead of linking information to genome coordinates. SubtiWiki also contains annotations of ncRNAs and UTRs, which are based on a large-scale study investigating the transcriptome of *B. subtilis* in over a hundred different environmental conditions using tiling-arrays^5^. However, SubtiWiki, DBTBS, and BsubCyc remain protein-centric^8, 20^, and predominately contain annotations of protein-coding genes and their functions. All of these resources annotate the tRNAs and rRNAs of *B. subtilis*. Oppositely, the annotation of less well-known ncRNAs have a lower quality in positional information and functional annotation. Rfam is a database of structured RNAs, genes or regulatory elements, from which covariance models have been made over homologs or families structured RNA^33^. These covariance models can be used to predict structure in genomic sequence^34^. Such screen in Bacillus Subtilis seem not to have been made and compiled into the annotation resources. Consequently we carried out such screen here. Yet additionally, there is still potential of discovering new bacterial ncRNA types and structures^35, 36^. Thus, an improved *B. subtilis* annotation would facilitate the discovery of coding and non-coding RNAs, their structures, and their regulatory functions^36, 37^. Moreover, an improved annotation could include the polycistronic relationship between genes and their expression, an aspect ignored in most bacterial RNA-seq analysis^38, 39^.

BsubCyc^31^, SubtiWiki^5, 8^, RefSeq^20^, literature references^5, 22^, and our own Rfam scan^33, 34^ provide a non-redundant genome annotation set, which fully contains comprehensive number of other resources listed in Table 1. Here, we describe the *Bacillus Subtilis* Genome atlas (BSGatlas), which improves upon the coordinate-based annotation of genomic elements covering coding and non-coding genes, UTRs, promoters, terminator sites, and operons by integrating existing resources and new genome-wide *in silico* predictions of ncRNAs. Using the information from the resources allows us to provide a more complete annotation of the *B. subtilis* genome, its genes, including ncRNAs, and to resolve inconsistencies between the available annotations. To facilitate the use of our annotation, we provide a UCSC browser based interface for visualization and a data download.

## Results and Discussion

### Enhanced annotation of coding genes, non-coding and structured RNAs

To obtain an enhanced and updated annotation of the genomic coordinates of coding and non-coding genes, we collected resource specific gene annotations for *Bacillus subtilis* strain 168 from BsubCyc^31^, SubtiWiki^5, 8^, RefSeq^20^, and the riboswitches found by Dar *et al.*^22^. In addition to these sources, we included our own computational screen of the *B. subtilis* genome for non-coding and structured RNAs using Infernal^34^ and *Rfam*^33^, because the most recent Rfam version has to our knowledge not been used in *B. subtilis* yet. We used the entire Rfam model collection, including models for non-bacterial RNAs, such that we had a negative control set while scanning at three different family-specific sensitivity levels: Conservative, medium, and sensitive. We deemed the most sensitive level to be too noisy, such that we excluded it from the gene annotation merging, yet we still make the annotation available in a separate genome browser hub (see gene annotation section in methods and results for genome browser hub) The screen resulted in a set of 214 conservative ncRNA matches, and additional 13 ncRNA at the medium level, which totals 227. Most of these *in silico* annotated ncRNAs (90.3%) are also in other resources (see below). 8 of these 13 ncRNAs were already annotated in RefSeq. The additional 5 ncRNA elements provided two *cspA* structures, which are temperature sensitive cis-regulatory structures^40^, the asRNA *rliD*^41^, the sRNA *bsrG*^42^, and an additional self-splicing intron^15^ within the ribonucleoside-diphosphate reductase gene *nrdFB*.

We also evaluated the annotation provided by SRPDB^28^, BSRD^27^. tmRDB^28^, tmRNA^29^, tRNAdb^30^, Ensembl^24^, PATRIC^25^, and IMG\M^26^. However, we did not include these as they are already contained in one of the earlier mentioned resources, based on an outdated RefSeq reference annotation, or are almost identical to it with respect to gene coordinates (trivial coordinate comparison not shown).

Before merging the annotations from the resources, we assigned to each of them a priority, which we use to resolve differences. When assigning these priorities, we consider the known context of the resources, such as technical limitations and dates of update. Our general guideline is to prefer manually curated over *in silico* annotations. To avoid mixture of coding and non-coding annotations and because we anticipate coding annotations to be more accurate in position (see below), we prioritize coding above non-coding resources. Among the listed resources, only BsubCyc and RefSeq annotate coding genes and their genome coordinates. We prioritize RefSeq over BsubCyc, because it contains a more recent expert curation of the *B. subtilis* genome coordinates^6^. For the annotation of non-coding and structured RNAs, we use both the remaining resources and the ncRNA parts from RefSeq and BsubCyc in a prioritized fashion. Overall, we prioritize literature-based information over predictions, with resolution constraint annotation having the lowest priority. We consider the coding and non-coding part for BsubCyc and RefSeq to be two separate sources each, such that we can assign them different priorities according to the above described approach. In Table S1 we summarized the individual priority values.

We investigated our assigned resource priorities by computing the Jaccard similarities of overlapping annotation pairs between and within the resources (Figure S1 and S2). These distributions of similarity values showed that nearly all overlaps between coding genes are either very similar (Jaccard *>* 0.98) or very dissimilar (*<* 0.1). Notably, the ncRNA genes had slightly lower similarities (*>* 0.9), and there were nearly no dissimilar ncRNA overlapping pairs. Coding sequences did not overlap ncRNAs except for riboswitches and a single ncRNA from a low confidence resource, which after manual inspection has been reclassified (see below). This similarity pattern was persistent across all resources and their assigned confidence levels. Yet, the ncRNA predictions made by Nicolas *et al.* during their tilling-array study tend to be less similar to the annotations from manual curated resources, which shows the limited in resolution due to the underlying technology. The limit in resolution combined with the biological constraint of nucleotide triplets on coding sequences confirms above stated assumption that the boundaries of of non-coding genes are less accurately annotated. We also investigated the distribution of Jaccard similarities for those gene pairs that correspond to each other according to matching gene names and locus tags. We found that all matching genes, both coding and non-coding, have a Jaccard similarity above 0.8, except for the spore coat protein *cotT* and three hypothetical proteins of unknown function (*yoyG*, *yqjU*, and *yrzH*). The genome positions of these four genes were largely divergent such that we decided to keep them for now as separate entries.

Based on the distribution of Jaccard similarities, we decided to merge two gene annotations if their similarity was at least 0.8, and for overlaps between two ncRNAs when their similarity was at least 0.5 or one of them is fully contained within the other. The special case of riboswitches overlapping coding genes was excluded from merging, allowing their annotation to be kept separate despite overlaps. Each group of genes, that fulfill the above criteria between each other, was merged into one. We used for the merging the coordinates that stemmed from the resource we have most confidence in, or if multiple resources had the same confidence level the union of the overlapping coordinates. The latter results in the longest stretch of annotation. Here, this only apply for the ncRNA resources, as the coding resources are consistent in the annotation in relation to the priority of the corresponding resources. because their boundaries are less clear (see assumption above) and trading-off between curated annotations might require non-trivial follow-up experiments. We assigned each merged gene the most specific gene type found in the group (eg sRNA instead of putative predicted ncRNA). Finally, we verified via locus tag and gene name matching that there were no erroneously merged genes or pair of genes that were missed. Please note, that in this approach we merge across all priority levels in a single step while at the same time allowing to merge annotation from the same level, possibly even within the same resource (see below).

Compared to the latest RefSeq assembly^20^, which is the standard reference for *B. subtilis* annotations^5, 8, 23, 37^, our merged gene annotation (Table 2) contained 229 additional ncRNA genes and structures, of which 61 ncRNAs have a clear annotation of the ncRNA type from literature resources (28) Rfam matches (23), or both Rfam and literature resources (10). In total, the additional 229 ncRNA genes originated predominately from literature resources (85) and the Nicolas *et al.* tilling-array study (112). Only 22 genes were uniquely described by Rfam, and 11 genes by Rfam and at least one of the other mentioned resources. The remaining ncRNAs of unknown type originated from the Nicolas *et al.* tiling-array study^5^. In total, the number of annotated known regulatory small RNAs, antisense RNAs, and riboswitches doubled in the merged gene set.

**Table 2.** Comparison of the resources with the merged results, aggregated by category, and how the merging process impacted the annotated gene types. The shown comparison statistics are relative to the gene annotation with the highest priority for each category. The full, un-aggregated table is in the supplementary Table S1.

|  | Merged gene annotation | RefSeq | BsubCyc | Literature resources | *Rfam* | Nicolas *et al.* predictions |
| --- | --- | --- | --- | --- | --- | --- |
| Protein Coding Genes | 4,332 | 4,324 | 4,188 | - | - | - |
| putative/predictions | 173 (4 %) | 88 (2 %) | 1,210 (29 %) | - | - | - |
| Hypothetical status removed | - | 9 (0 %) | 1,204 (29 %) | - | - | - |
| Resource specific genes | - | 144 (3 %) | 8 (0 %) | - | - | - |
| Non-Coding RNAs | **441** | **212** | 183 | 91 | 227 | 196 |
| ribosomal RNA | 30 (7 %) | 30 (14 %) | 30 (16 %) | 0 (0 %) | 30 (13 %) | 0 (0 %) |
| transfer RNA | 86 (20 %) | 86 (41 %) | 86 (47 %) | 0 (0 %) | 86 (38 %) | 0 (0 %) |
| small regulatory RNA | **36** (8 %) | **14** (7 %) | 9 (5 %) | 0 (0 %) | 30 (13 %) | 0 (0 %) |
| regulatory antisense RNA | **7** (2 %) | **3** (1 %) | 2 (1 %) | 0 (0 %) | 3 (1 %) | 0 (0 %) |
| riboswitch | **103** (23 %) | **55** (26 %) | 26 (14 %) | 82 (90 %) | 72 (32 %) | 0 (0 %) |
| self-splicing intron | 3 (1 %) | 0 (0 %) | 0 (0 %) | 0 (0 %) | 3 (1 %) | 0 (0 %) |
| ribozyme, SRP, tmRNA | 3 (1 %) | 2 (1 %) | 2 (1 %) | 0 (0 %) | 3 (1 %) | 0 (0 %) |
| putative/predictions | 173 (39 %) | 22 (10 %) | 28 (15 %) | 9 (10 %) | 0 (0 %) | 196 (100 %) |
| Hypothetical status removed | - | 18 (8 %) | 29 (16 %) | 10 (11 %) | 0 (0 %) | 26 (13 %) |
| Reclassified as coding | - | - | - | - | - | 2 |
| Resource specific genes | - | 7 (3 %) | 1 (1 %) | 24 (26 %) | 22 (10 %) | 170 (87 %) |

We investigated the refinement of coordinates in our merged gene annotation with respect to both absolute and relative changes (Table S2 and Figure S4). Because RefSeq’s coding genes were assigned the highest priority in the merging step, no coordinates changed for them. The only peculiar exception is the merging of a predicted open reading frame with the annotated coding gene *cotG*, which fully contained the prediction. Upon closer inspection, we found in the corresponding RefSeq entry (BSU36079) the comment “doubtful CDS”, such that we assume this to be an artifact we resolved. We observed that 49 coding genes annotated in BsubCyc had coordinate changes between 10 and 250 bp, although due to the lengths of genes the overall Jaccard similarity between the respective original annotations and the merged gene annotations is barely reduced. We mentioned above 4 highly divergently annotated coding sequences with low Jaccard similarity between RefSeq and BsubCyc. Besides these, BsubCyc only uniquely annotated three more hypothetical proteins (*yfmA*, *ypzE*, and *ytzK*). Otherwise, all coding genes of BsubCyc were contained in RefSeq. Thus, our merging had a limited effect on gene coordinates, likely reflecting the high quality of the protein annotation. In contrast, 70% of RefSeq’s and 79% of BsubCyc’s annotated non-coding genes and structures coordinates were refined up to 500 bp, although even smaller changes of 50 to 100 bp reduced the Jaccard similarity down to 0.6 or more. These changes of BsubCyc’s and RefSeq’s non-coding annotation were mainly the result of the merging with higher confidence resources from the literature-based resources. Moreover, two putative ncRNA from the Nicolas *et al.* tiling-array study (S458 and S1078) were reannotated as part of the three-component toxin-antitoxin-antitoxin system *SpoIISABC* (BSU12815) and as a separate hypothetical short peptide (BSU28509).

After the gene merging procedure, we associated the meta-information, including general descriptive texts, synonyms, molecular weights of translated proteins, and literature references, with the merged gene annotations. In addition, we added BsubCyc’s functional annotation with Gene Ontology terms^32^ for 71% of all genes and Enzyme Classifications from SubtiWiki^8^ to 14.5% of all genes, RefSeq^20^ to 19.5%, and BsubCyc^31^ to 15.7%, which in combination covers nearly 22% of genes. We recoded these Enzyme Classifications into a human-readable form according to the definitions in the BRENDA database^43^. Finally, we added the information available in SubtiWiki’s *B. subtilis* specific category system^8^, which contained information about 91.9% of genes. From the Nicolas *et al.* tilling-array study^5^, we listed experimental conditions with highest and lowest expression and in expression correlated genes. We did not further filter the meta-information, but for each piece of information we indicate its origin.

### Promoter map and complex operon architecture

Bacterial genes are often co-expressed in operons, defined as a set of genes that can be transcribed into a single RNA transcript from one promoter region^16, 18^. Even with the absence of splicing, bacterial RNA expression can be quite complex, because of the use of alternative promoters and transcription termination^16, 44^ (illustrated for the *alaS* operon in Figure 3d). Only few resources annotate full transcripts. More commonly, transcriptional units (TUs) annotate sets of genes that can be transcribed together without exact indication of transcript boundaries. The combination of multiple resources described a set of 2, 473 TUs.

We collected the TSS and TTS annotations from the external resources and determined their resolution limit by investigating the distribution of distances between annotations (Figure S6). The manually curated annotation resources, BsubCyc and DBTBS, had a single nucleotide resolution on their TSSs, whereas the TSSs predicted in the Nicolas *et al.* tiling-array study had a resolution window of 50 bp, which is about twice the probe length of the underlying tiling-array design^5, 45^. Additionally, the tiling-array study provided as 5’ UTR classified transcribed regions that were without a clear TSS signal. Thus, these positions should resemble a TSS with reduced accuracy. Accordingly, we found their resolution windows to be larger, about three times the probe length. We found similar resolutions windows per resource with respect to TTSs. Using a merging similar to the one used for genes (see methods on TSS/TTS map), were retrieved a unified set of 3, 397 TSSs and 2, 666 TTSs (Figure 1). The intersection of annotations from BsubCyc and Nicolas *et al.* The underlying annotation resources shared a set of 12.5% of the TSSs and 23.7% of the TTSs. The largest quantity of annotation originated from the Nicolas *et al.* tiling-array study^5^. In the combined set, 79.2% of the TSSs and 57.6% of the TTSs are solely provided by the tiling-array data. In contrast, the union of the higher resolution resources BsubCyc and DBTBS provided unique annotations for 3.4% of TSSs and 17.3% of TTS. Overall, a total of 19.4% of the TSSs had both single-nucleotide resolution and en expert curation origin. For the TTSs the proportion is 41.0%. In comparison, the distribution of sigma factor binding sites associated with the TSSs remained similar, yet not as many promoters have an unknown sigma factor as in BsubCyc.

**Figure 1.**
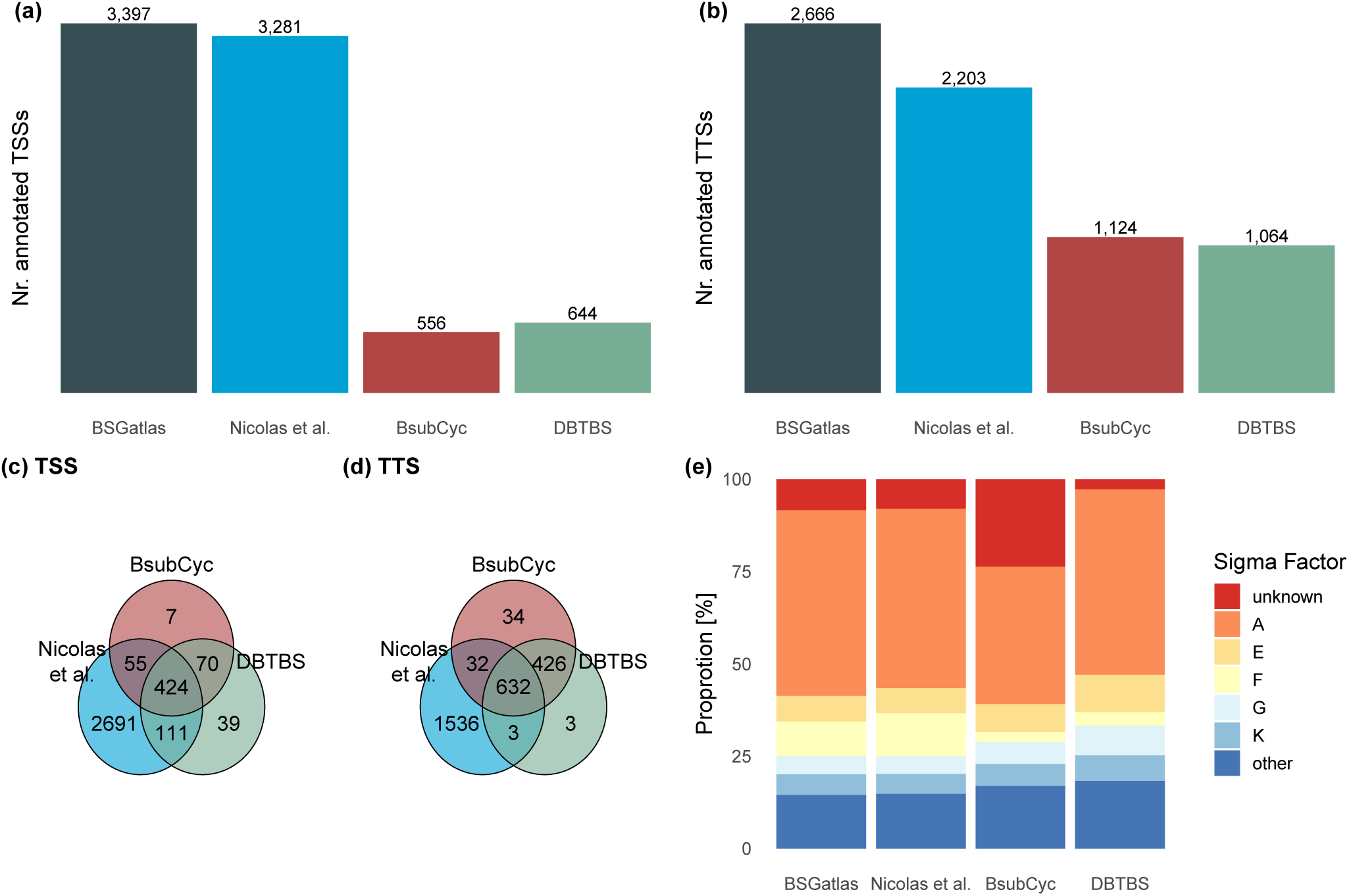
(a) Comparison of total numbers of TSSs and (b) TTSs provided by the individual resources and the unified set in our BSGatlas. Venn diagram how many (C) TSSs and (d) TTSs are shared between the resources. (e) Comparison of proportions of which sigma factor binding sites are annotated in the resources.

The UTRs annotated by Nicolas *et al.* were divided into non-overlapping pieces. Yet, the sum of their lengths with the distances to the associated genes show that the biological 5’ and 3’ UTRs in *B. subtilis* are at most 2, 000 bp long (Figure S7). From our unified TSS annotation, 94.6% of all TSSs are within that distance to the 5*^!^* end of upstream coding or non-coding genes in the merged gene annotation, and 95.5% of TTS to the 3*^!^* ends of downstream genes. (Table S3 and Figure S7). Thus, the regions from a TSSs to upstream genes, within the 2, 000 bp distance cut-off, represent 5’ UTRs and respectively 3’ UTRs. After filtering for a minimal length of 15 bp, we derived a map of 5’ UTRs (2, 943) and 3’ UTRs (2, 095). This approach associated 5’ UTRs (103) to upstream non-coding RNA structures; such that the UTRs fully reflect the biological untranslated region, we extended these up to the start of the next upstream coding gene.

Finally, we inferred 1, 126 internal UTRs from the regions between genes that are known to be co-transcribed according to the above-described combined TU list. For UTR lengths higher than 46bp the distribution of our *in silico* obtained UTRs lengths are similiar to those of the ones obtained by (*p_i_ <* 7.93 *·* 10*^−^*^8^ two-sample Kolmogorov-Smirnov test for all UTR types *i*) (Figure S7). Below the tiling array resolution limit of 50bp, the Nicolas *et al.* data can not provide UTR annotations, whereas we found 2, 343 UTRs as short as 15 bp.

We compared our predicted UTRs with those found by Nicolas *et al.* by identifying pairs that overlap in at least 25 bp and compare their types (Table S4). Overall, 93.3% of their 5’ and 3’ UTRs agreed with our computed UTRs. We manually inspected the 56 remaining 5’/3’ UTRs on why we do not directly annotated these. The Nicolas *et al.* supplement itself classified 24 as curated independently transcribed ncRNA genes. One was characterized as the sRNA *asrE* by the updated RefSeq annotation. Due to our TSS and TTS improvement in resolution, we did not see the basis for 3 UTRs. The *>* 500 bp long 5’UTR S422 is almost fully contained by an anti-sense operon, such that the classification as UTR could be discussed. 17 overlap in at least 50 bp with a gene in the merged gene annotation, and thus were indeed translated regions. 10 UTRs were longer than 2000 bp up to more than 4000 bp, such these are not within our expectation of UTR lengths, such that these should be investigated more closely. Only 74.2% of the Nicolas *et al.* internal UTRs were in agreement with our findings. By considering available TUs, we found that *∼* 7% of the internal UTRs could be separated into distinct 5’ and 3’ UTR elements. Furthermore, we observed that 47% of what Nicolas *et al.* classify as transcribed intergenic regions overlap in at least 25 bp with our UTRs. Thus, we deem these not to be biological intergenic regions. We suspect two main reasons for these misclassifications. On the one hand, the tilling-array study had a resolution limitation, which leads to difficulties in identifying separate UTR elements, such as described above. On the other hand, the TU annotation sources—that we used—had not been published and thus were not available at that time. Yet, even with the availability of higher-resolution RNA-seq data, the lack of TU annotation hinders the prediction of internal UTRs and makes it a non-trivial endeavor^46^. The overlap comparison showed that 76.4% of our computed UTRs had no correspondence, and thus these might be new UTR annotations instead of refinements. In total, we annotated five times more UTRs than the Nicolas *et al.* annotations.

We used the associations we found between TSSs, TTSs, UTRs, and genes with respect to TUs to computed transcripts (Figure S3 c). By following TSSs or TTSs that were inside of annotated TUs, we inferred 815 novel TUs. In total, we found 4, 804 transcripts. The number of transcripts is larger than those of TUs, because a TU can have multiple transcripts with varying UTR lengths. In comparison, BsubCyc annotated only 1, 602 transcripts. We computed a set of 2, 309 operons, based on the criteria that two transcripts that share the full sequence at least one gene are isoforms from the same operon (Figure S3 d). For 99.7% of these operons, we annotate at least one transcript which contains all of an operon’s genes. Afterwards, we looked up for each transcript which TU annotation it was based on, and from which resource the TUs stemmed. Subsequently we computed the proportion of genes for which each resource provide transcriptional annotation. The unified set annotated transcripts for 84% of the genes, which is 3 percent points more than the largest individual resource. However, combined with our novel TUs nearly 93% of all genes have a transcript annotation.

Finally, we investigated how the distribution of our computed operons compare to related micro-organisms. We classified our operons similar to the description from recent studies focused on operon annotation in *Streptococcus pneumoniae*^47^, and *Escherichia coli*^18^ (Figure 2). Of our operons without any isoforms, 42.8% are simple, monocistronic operons and only 9.1% are traditional, polycistronic. A total 45.1% are complex operons, meaning that they have alternative isoforms due to alternative transcription start and termination sites. In comparison, *E. coli* has 45% simple, 19% traditional, and 36% complex operons^18^. For *S. pneumoniae* the distribution is 47% simple, 10% traditional, and 43% complex operons^47^. We computed the distribution of the number of genes, internal TSS, and internal terminator per operons. We observed visually similar distributions as *S. pneumoniae* (Figure S5 compared to Warrier *et al.*’s Figure 3B^47^), although we annotated more monocistronic operons without known TSSs or TSSs. Given that our inference is based on already curated TUs, we claim that our operon annotations are biologically reliable. However, we showed that the distribution of operon classes are comparable to other microorganisms, which further supports that statement. A homology based comparison of the annotations between the organisms would be needed for a definite statement about the phylogenetic conservation, yet, this would be outside of the scope of this annotation effort.

**Figure 2.**
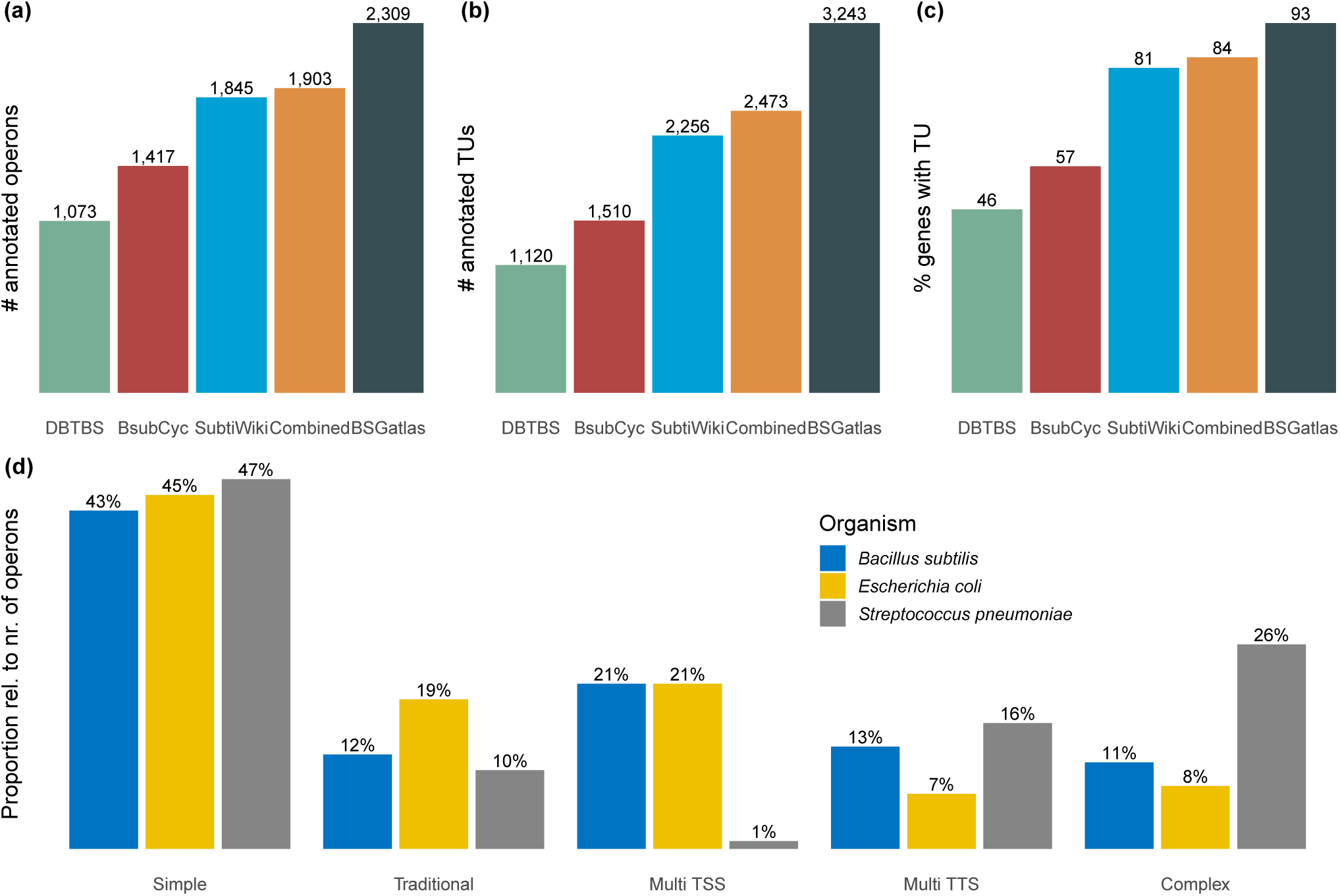
Comparison of the number of (a) operons and (b) TUs the individual resources provide, and (c) what proportion of the merged gene set they cover. (d) Comparison of the operon classes for our computed operons in *B. subtilis* with those found in *S. pneumoniae* and *E. coli*^18, 47^.

### Genome browser hub with enhanced information access

The UCSC Genome Browser provides the means to easily visualize and share track information from sequencing experiments and annotation sources^48^. There is a *B. subtilis* genome browser in UCSC’s archaeal genome section^49^, yet this hub has the same protein-centric focus as discussed earlier and has a quite limited set of data tracks. Thus, we compiled our BSGatlas annotation as an assembly hub (Figure 3d), thereby allowing users to investigate their data side by side with our annotation, without the need to install dedicated software on their machines. Because of the powerful feature set of the UCSC browser framework, the type of data a user can visualize is plentiful: These could be large-scale high-throughput coverage data, custom annotation tracks, or even sequence alignments. We enabled the browser hub to search for all genes by their names, synonyms, and loci identifiers, including the alternatives and spelling variants available from all used resources. For each annotated gene, transcript, TSS, etc., we provide a detailed summary page that contains all meta-information that we retrieved from externally available resources (Figure 3c). For each piece of meta-information, we indicate from which of the external resources it originated, and we link back to the external resources. Moreover, our browser contains the tracks for the resources used to create BSGastlas, meaning that users can compare the original annotation themselves. The browser also includes a table browser and a BLAT search option^50^, which facilitates easy download of data set and identification of *B. subtilis* genomic sequences, respectively. To ease the navigation of the large number of annotation tracks the BSGatlas provides, we grouped them. A user can interactively activate the display of groups in the browser, and they can select for each group individually their tracks of interest (Figure 3e).

**Figure 3.**
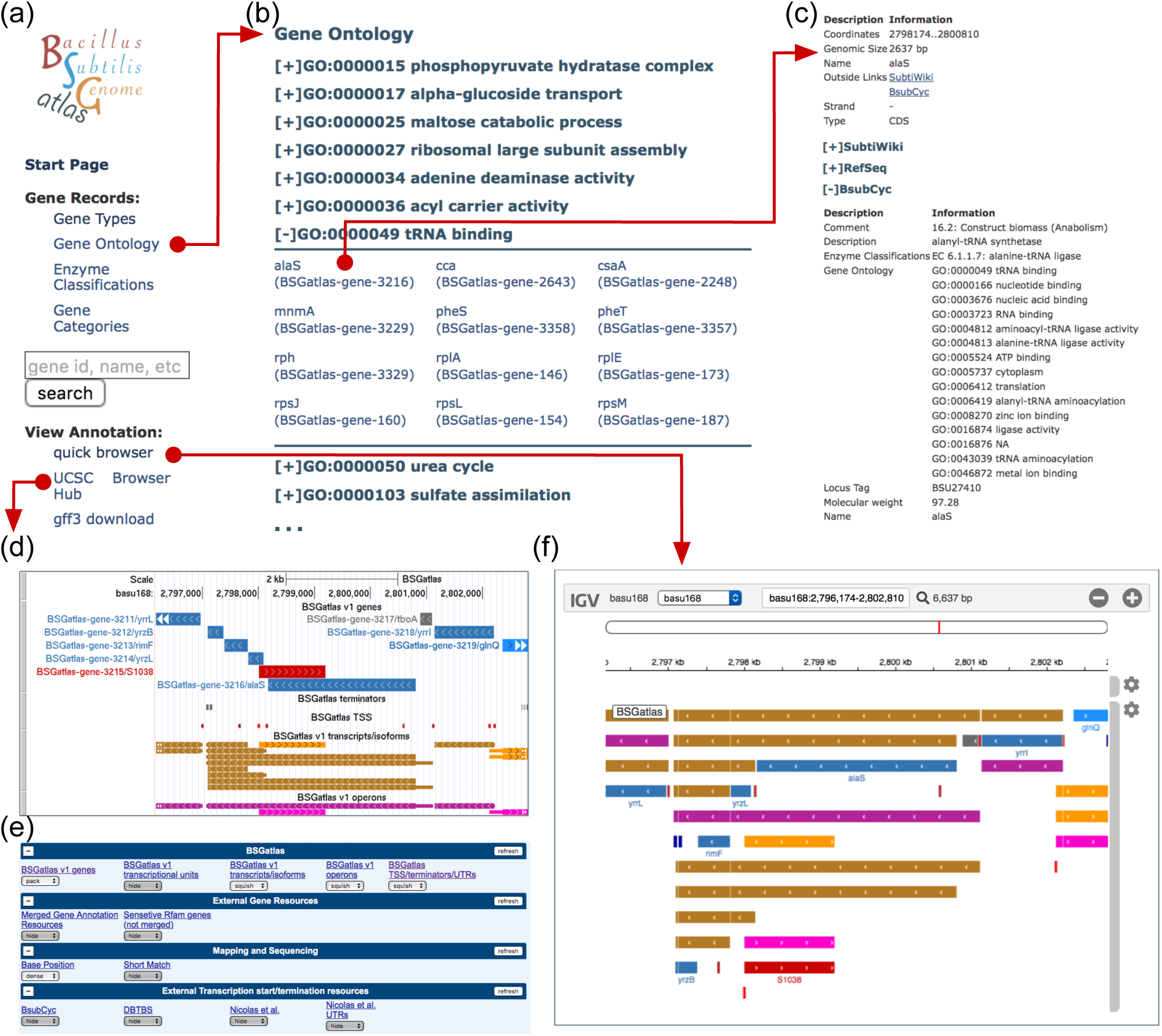
Illustration of the BSGatlas and its features. (a) The BSGatlas start page provides the main groups of navigational entries that lead either to the gene records, the search function, or the various visualization options. (b) The gene record pages have the same structure across each classification system, that are gene types, enzyme classes, or functional classification as listed in SubtiWiki or Gene Ontology terms. Each group is shown as a clickable entry, which list its associated genes. The gene names are shown as links to (c) detailed description pages. These show all meta-information we found for all genes and all other annotated entries, such as transcripts and operons. (d) UCSC browser. The user has the option to directly show the BSGatlas annotation as an assembly hub in the UCSC browser. Thus, they can also show their data, eg an RNA-seq experiment, right next to the annotation. (e) UCSC browser control. Underneath the UCSC browser panel, a user can control details of what parts of the BSGatlas are shown. This includes also the gene coordinates as they were originally annotated, which allows a more closely investigation of the gene merging process. (f) Quick browser. We provide a fast visualization directly in BSGatlas main page, which allows a user to get a quick overview of the annotation without the need to leave the webpage. A click on any BSGatlas annotation redirects to the corresponding description page.

On each detailed description page, we included a lightweight browser (based on the *igv.js* library^51^) (Figure 3f), which allows a user to get a fast overview of the genomic context of an annotation. We also provide the annotation as a GFF3 download option (Figure S3e) to facilitate offline visualization of the BSGatlas with programs such as IGV or IGB^51, 52^. We use a unified color scheme across the different visualization options. The color scheme indicates the gene types (protein, tRNA, rRNA, etc) and the strand location (Figure S8). Additionally, we provide gene record information for the various gene types and gene set; these are at the moment enzyme classifications, functional annotation with GO terms, and SubtiWiki’s category system (Figure 3a+b)). Thus, the BSGatlas now offers access to gene records in a single combined resource, and users can compare the meta-information between the resources more easily. The BSGatlas can be accessed under: https://rth.dk/resources/bsgatlas/

## Conclusion

*Bacillus* species are important for cell-based protein production and *B. subtilis* is probably the most studied Gram-positive bacteria. Thus improvements in the quality of its genome annotation have scientific added value.

In this work, we carefully integrated the existing annotations from BsubCyc^31^, SubtiWiki^5, 8^, RefSeq^20^, literature references^5, 22^, and our own Rfam scan^33, 34^ which resulted in a single atlas annotation for *B. subtilis* that comprises genes, transcripts, and operons. In addition, we updated the Rfam annotation of the *B. subtilis* genome. In combination, this led to an increase in the number of non-coding genes by 229, which stemmed from literature resources (85) and the Nicolas *et al.*, tilling-array study (112), and Rfam (33, with 11 genes being also contained in the other resources). We inferred a unified set of 3, 397 TSSs and 2, 666 TTSs Overall, the external resources indicate a high curation level and resolution confidence in 41% of all TTSs. The annotation of TSSs still lack substantially behind, as existing curation provide a high-resolution for only 19.4% of TSSs. Yet, these percentages were only achieved by integrating existing annotations, which is an improvement over currently available data. For each annotated gene and ncRNA, we carefully collected meta-information, thereby allowing researchers to find the currently available information in one location. Importantly, we also provide links to the original data sources for easy access. Our new annotation, and the data sources used for making it, are available in the UCSC browser format. In this way *B. subtilis* researchers can easily visualize and download data as well as compare to their own data.

About half of all TTSs had expert curation. Although our TSSs unification brought about some improvement, the quality of TSSs still lacked behind. Furthermore, we curated a list of TUs, which in combination with the TSSs and TTSs allowed us to predict UTRs and infer novel TUs. This combined annotation gave rise to a complex architecture of operons and their isoforms. The distribution of operon features and the operon classification were similar to those described in other Bacteria. In total, we curated the most comprehensive annotation of coding and non-coding genes in *B. subtilis* to date, and we provide transcriptional annotation for nearly 93% of all genes. We compiled our annotation as an interactive online annotation browser. Also, we put the annotation in a standardized format for usage in various-omics studies.

For *B. subtilis*, our bacterial genome and transcriptome annotation is to our knowledge most comprehensive of its kind. Thus, we anticipate that it will complement existing resources and be in studying the function of the genome including non-coding genes and in considering transcriptional relationships of genes in transcriptomic studies. Another use case could be to use our annotation as a reference in bench-marking transcript and operon prediction methods. On a statistical level, we anticipate that our annotation could be used to model co-transcription and the implied statistical dependencies in differential expression analyses, an aspect commonly ignored at the cost of reduced statistical power. As a consequence of this integrative annotation, we enriched the genome annotation of *B. subtilis* with a comprehensive list of TSSs and TTSs, operons and their isoforms, and an unprecedented level of untranslated region (UTR) annotations, and RNA structure predictions.

## Methods

### Notes on general computational workflow

All analyses, if not otherwise indicated, were performed in `R` 3.5.2^53^. We utilized a multitude of `Bioconductor` packages^54^ and the `tidyverse` collection^55^. The predominately utilized packages were `rtracklayer`^56^, the annotation packages `GenomicRanges`^57^ and `plyranges`^58^, the parser `genbankr`^59^, the color palette `ggsci`^60^, the library for nucleotide sequence handling `Biostrings`^61^, the graph analysis tool `tidygraph`^62^, and the table creation package `kableExtra`^63^. For improved reproducibility of the annotation construction, all steps were conducted in an Anaconda environment. Thus, the exact list of versions for these packages and all of their dependencies at each step of the annotation creation is explicitly stated. The anaconda environment and the scripts including intermediate computational results are available at doi 10.5281/zenodo.3478329.

### Gene annotation resources

We built the annotation according to the latest RefSeq reference assembly (accession `ASM904v1`^20^), which contained the major gene annotation refinement from February 2018^6^. We used this annotation as described in the *GenBank* file. Based on the provided human-readable gene description text, we were able to determine the specific non-coding RNA type for 92% of the 212 non-coding genes.

The BsubCyc database^31^ version 38 (released Aug 9, 2017) has a systems view representation. We built a custom parser to extract for BsubCyc’s database, which contains 4, 188 coding and 184 non-coding genes and structures. Also, its curation contained experimentally verified information about 574 TSSs with nucleotide resolution and 1, 246 TTSs. These provide clear transcribed boundaries for some of the 1, 602 TUs from BsubCyc, yet explicit UTR elements are note annotated. As meta-information, it provides GO-terms and indication of transcriptional regulatory relationships. In addition, BsubCyc gives detailed metabolic and enzymatic reaction and pathway overview, which will not be considered here.

The genome was scanned for the 3, 016 families of *Rfam* 14.1^33^ using *Infernal’s cmsearch* version 1.1.1^34^. The scan was conducted with a family-specific score cutoff matching half the so-called gathering score. The hits from the scan are reported at three sensitivity levels. (1) At the conservative level, an E-value cut-off at 10*^−^*^6^ and a match score of at least the gathering score was applied; (2) at the medium level, only the E-value cut-off at 10*^−^*^6^ was used; and (3) at the sensitive confidence level, the E-value requirement was relaxed to 10*^−^*^3^. The results were post-filtered using an updated version of the RNAnnotator pipeline^64^. Within each level of confidence all hits overlapping at least 40 bp were merged and the best hit by E-value was chosen (or by score if E-values were tied). At the respective sensitivity levels, the scan identified 214, 227, and 265 non-coding genes and structures. We did not filter the Rfam collection before the scan, such that apparent false-positive cases are included. We used these to determine the sensitivity to be used in the gene merging. After inspection, we decided against using our highest sensitivity level of *Rfam* for two reasons: (1) Due to the erroneous prediction of eukaryotic-specific ncRNA families, and (2) the too large number of statistically expected random predictions, which is given by the sum of E-values per sensitivity level (conservative = 9.52 *·* 10*^−^*^8^, medium = 9.88 *·* 10*^−^*^7^, sensitive = 0.0141).

The annotation specific to this highest sensitivity level is still shown in the browser.

Nicolas *et al.*^5^ conducted a large scale tiling-array study in over one hundred different environmental conditions. Within the resolution limit of the tiling-array, they identified 3, 242 TSSs and 2, 126 TTSs. Under consideration of these TSSs and TTSs, they identified 1, 583 novel transcribed regions. Depending on the location, they classified these as 1, 430 UTRs and 153 novel transcripts. Their supplementary material includes a literature survey, which provided the status of experimental verification for 81 ncRNA predictions. Furthermore, it has 9 additional ncRNAs that were not found by their tiling-array study. Dar *et al.*^22^ found a set of 82 riboswitches that they found by investigating transcription termination patterns in a term-seq experiment.

### Merging of gene annotations

We assigned to each resource a priority. We compared each overlapping pair of genes, and computed for each overlapping pair the Jaccard similarity, which applied for annotations is^65^

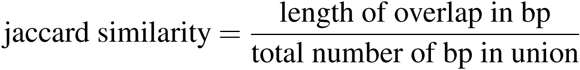

After inspection, we assume identity between an overlapping pair if it is not between riboswitches and coding sequences, and their Jaccard similarity is at least 0.80, or for overlaps between two ncRNAs of at least 0.5 or the special cases that one annotation is fully contained within the other. We identified all pairs of genes that fulfill this identity criteria. Assuming transitivity, we select groups of genes that are identical via computation of components in a graph. For each group, we collect the coordinates from the resource from the highest priority, or the union if there are multiple genes of equal priority (Figure S3a). A special case of gene merging could be the fusion of annotation from the same confidence level, and even from the same resource.

Afterward, we identified for each group the most specific type description across all priorities. In most cases, these were the type which is not putative. In only few cases we resolved ambiguities by preferring the type description asRNA over sRNA, and sRNA over riboswitch. The latter ambiguity is caused by the term-seq resource, which did not investigate possible additional biological functions of partially transcribed RNAs.

### Parsing DBTBS

Upon request, the authors of DBTBS^23^ kindly provided us with their latest annotation in an XML format. Unfortunately, this file did not contain coordinates for the most recent *B. subtilis* 168 reference genome sequence. Instead, it stated only nucleotide sequences for a seemingly outdated genome assembly. We utilized an exact sequence lookup to find coordinates for 98% of the 1, 262 annotated transcription factor binding sites and for 90% of the 1, 031 annotated TTS. To reduce erroneous or ambiguous annotations, we used unique matches without allowed mismatches. From the sigma factor binding sites that have the relative position to the TSS provided, we were able to infer a set of 644 high-resolution TSS positions.

DBTBS had a curated annotation of operons for 2, 201 genes. For 98.9% of these genes, we were able to find the corresponding gene in our merged set by comparing the gene names and keeping only unambiguous matches. Due to the high matching success rate of the genes, we were able to fully restore coordinates for almost all (98.6%) of the annotated 1, 123 operons.

### Parsing SubtiWiki

SubtiWiki provided a magnitude of meta-information, such as its gene categorization and lists of transcriptional regulations and interactions^8^. In the most recent version, SubtiWiki included some parts of BsubCyc^8, 31^, yet the curated list of TSSs, TTSs, and functional annotation via GO terms were not. Unfortunately, SubtiWiki does not provide the export of gene coordinates, such that we restored these from our merged gene set via comparison of locus, gene names, and synonyms. We were able to infer positions for 99.7% of the 5, 999 coding/non-coding genes, structures, and untranslated regions described in SubtiWiki; for 99.6% of the 2, 267 provided SubtiWiki TUs.

### Transcriptional Units

We collected a set of 1,602 (TUs) from BsubCyc^31^, 2, 259 TUs from SubtiWiki^8^, and 1, 123 TUs that are implied by operons from DBTBS^23^.

We investigated the overlaps of these TUs with the merged gene set and found two main overlap scenarios: Either genes were fully contained by a TU, or a gene or structure has a large overlap with a TU, possibly to the extent that it fully contains it. The latter was the case for a small peptide sequence that is fully contained in a known small regulatory RNA. Thus, we decided to add a gene or structure to a TU if (a) it is fully contained by the TU, (b) if it contains the TU, (c) or the overlap relative to the length of the gene is at least 70%. Overall, we added genes to 46 TUs from BsubCyc, 753 TUs of DBTBS, and 1,968 TUs from SubtiWiki. We removed 2 TUs, one from DBTBS and SubtiWiki each, which were erroneous as they would span more than a quarter of the whole genome. The resulting set of unified TUs totaled 2, 474 TUs.

### TSS and TTS map

We unified existing annotations of promoters and their TSS and TTS. We followed, for each type separately, an approach similar to the gene merging step. We first compared the distances and overlaps between the annotation to determine their resolution limits (Figure S6). We determined for each data type the resolution limits (see results). This resolution limit is the minimal distance to differentiate two separate annotations within a single resource. Thus, the true positions are up to half the resolution limit up- or down-stream of the annotated position. Here, we assume that annotations that overlap on the interval of the possible true positions might correspond to each other. A group of annotations that might correspond to each can only contain two annotations of the same resolution if there is also an annotation with worse resolution in the group. Consequentially, we could retrieve a unified TSS and TTS map by finding these group with a graph-based method similar to the gene merging, and then subsequently taking per the annotations that had a resolution equal to the minimal group resolution. For each combined set of TSSs, we collected the information which sigma factor promotes the transcription from the entry with the lowest resolution limit.

### Untranslated regions

Within a distance cut-off of 2, 000 bp, we created a TSS and TTS map by associating TSS to the gene with the closest 5’ end and TTS to the one with the closest 3’ end (Figure S7 and S3b). If there was a gap between a TTS and the associated gene of at least 15 bp length, we created a 3’ UTR in its place. We similarly created 5’ UTRs based on TSSs. However, we extended the length of 5’ UTRs up to the next coding gene if the first direct association is a non-coding RNA structure. We filled gaps longer than 15 bp between the genes listed in the TU with internal UTRs. The only exception is that we did not add an internal UTR for the *sigK* TU, because the over 10, 000 region within it is not actually transcribed due to its unique regulatory mechanism^66^.

### Operon architecture

We inferred novel TUs and the full transcripts by a graph-based approach (Figure S3c). We created a direct graph that contains the transcriptional relationship and directions of TSSs, TTSs, UTRs, and genes. We added dummy TSS and TTS for TUs whose genes had none associated, as computed earlier. For all paths that connect TSS and TTS, we created a transcript that contains the genes and UTRs along each path. We created new TUs for those paths and transcript that contain a set of genes not contained in our TU list.

Afterward, we computed operons from these transcripts, by finding connected components in a graph. The graph contained nodes for each transcript and each gene; an edge was added between them if the transcript contains a gene (Figure S3d). Due to the particularity of the *sigK* transcriptional regulation^66^, we excluded the associated TU both from the transcript and operon inference.

### Browser hub

We generated the browser hub according to the official UCSC browser documentation and converted the tracks into the custom binary format with the UCSC tools. The individual tracks were organized according to the track definition (https://genome.ucsc.edu/goldenPath/help/trackDb/trackDbHub.html) and the search functionality was provided as a Trix index (https://genome.ucsc.edu/goldenPath/help/trix.html).

## Acknowledgments

This work was supported by the Innovation Fund Denmark [5163-00010B]. The authors want to acknowledge Betina Wingreen Jensen for project coordination, Yuko Makita for kindly providing the DBTBS data, and the BSubCyc and SubtiWiki maintainers for providing direct downloads.

## Author contributions

CA and SES conducted the Rfam scan. LDP and JV validated the atlas annotation. CA and ASG compiled the browser hub. EGT was involved in the discussion of methodology, results, and structure of the manuscript. TKK sanitized the user front-end. Also, EGT and JV reviewed the entire manuscript. ASG designed and conducted all other analyses and wrote the manuscript. JG coordinated the study and reviewed the manuscript.

---

**Table S1.** Comparison of the individual gene annotation resources with the merged gene set. The topmost row contains the priority we assigned to that source, with a numerically lower value indicating higher confidence. Our general guideline behind the assigned confidence levels were to give higher priority (i) to resources annotating protein-coding genes, in order to avoid confusion of the overall clear boundaries of coding genes with the less clear boundaries of non-coding genes or structures, (ii) to the more recent resource, (iii) and to prefer expert curated or literature review resources over computational ones. We considered resources with equal priorities to be equally trustworthy, such that we joined their annotations with the union of the coordinates.

|  | Merged gene annotation | RefSeq Coding | BsubCyc Coding | RefSeq Non-Coding | Nicolas *et al.*'s literature review | Nicolas *et al.* trusted predictions | *Rfam* (conservative) | BsubCyc Non-Coding | Dar *et al.* term-seq | *Rfam* (medium) | Nicolas *et al.* predictions |
| --- | --- | --- | --- | --- | --- | --- | --- | --- | --- | --- | --- |
| Resource Priority | - | 1 | 2 | 3 | 3 | 3 | 3 | 4 | 4 | 5 | 5 |
| Protein Coding Genes | 4,332 | 4,325 | 4,188 | - | - | - | - | - | - | - | - |
| putative/predictions | 173 (4 %) | 88 (2 %) | 1,210 (29 %) | - | - | - | - | - | - | - | - |
| Hypothetical status removed | - | 9 (0 %) | 1,204 (29 %) | - | - | - | - | - | - | - | - |
| Resource specific genes | - | 144 (3 %) | 8 (0 %) | - | - | - | - | - | - | - | - |
| Non-Coding RNAs | 441 | - | - | 212 | 9 | 81 | 214 | 183 | 82 | 227 | 115 |
| ribosomal RNA | 30 (7 %) | - | - | 30 (14 %) | 0 (0 %) | 0 (0 %) | 30 (14 %) | 30 (16 %) | 0 (0 %) | 30 (13 %) | 0 (0 %) |
| transfer RNA | 86 (20 %) | - | - | 86 (41 %) | 0 (0 %) | 0 (0 %) | 86 (40 %) | 86 (47 %) | 0 (0 %) | 86 (38 %) | 0 (0 %) |
| small regulatory RNA | 36 (8 %) | - | - | 14 (7 %) | 0 (0 %) | 0 (0 %) | 29 (14 %) | 9 (5 %) | 0 (0 %) | 30 (13 %) | 0 (0 %) |
| regulatory antisense RNA | 7 (2 %) | - | - | 3 (1 %) | 0 (0 %) | 0 (0 %) | 2 (1 %) | 2 (1 %) | 0 (0 %) | 3 (1 %) | 0 (0 %) |
| riboswitch | 103 (23 %) | - | - | 55 (26 %) | 0 (0 %) | 0 (0 %) | 63 (29 %) | 26 (14 %) | 82 (100 %) | 72 (32 %) | 0 (0 %) |
| self-splicing intron | 3 (1 %) | - | - | 0 (0 %) | 0 (0 %) | 0 (0 %) | 1 (0 %) | 0 (0 %) | 0 (0 %) | 3 (1 %) | 0 (0 %) |
| ribozyme, SRP, tmRNA | 3 (1 %) | - | - | 2 (1 %) | 0 (0 %) | 0 (0 %) | 3 (1 %) | 2 (1 %) | 0 (0 %) | 3 (1 %) | 0 (0 %) |
| putative/predictions | 173 (39 %) | - | - | 22 (10 %) | 9 (100 %) | 81 (100 %) | 0 (0 %) | 28 (15 %) | 0 (0 %) | 0 (0 %) | 115 (100 %) |
| Hypothetical status removed | - | - | - | 18 (8 %) | 9 (100 %) | 22 (27 %) | 0 (0 %) | 29 (16 %) | 1 (1 %) | 0 (0 %) | 4 (3 %) |
| Reclassified as coding | - | - | - | - | - | 1 | - | - | - | - | 1 |
| Resource specific genes | - | - | - | 7 (3 %) | - | 58 (72 %) | - | 1 (1 %) | 24 (29 %) | 5 (2 %) | 112 (97 %) |

**Table S2.** Comparison of the coordinates from each gene annotation resource with the coordinates of the resulting genes after merging. Shown are the number and the amount of refinements per annotation resource. The comparison under consideration of the gene lengths is shown in Figure S4.

| Resource | No difference | [1,10] | (10,50] | (50,100] | (100,250] | (250,500] |
| --- | --- | --- | --- | --- | --- | --- |
| RefSeq Coding (0) | 4,324 | 0 | 0 | 0 | 1 | 0 |
| BsubCyc Coding (1) | 4,139 | 18 | 13 | 12 | 6 | 0 |
| Nicolas *et al.* trusted predictions (2) | 71 | 3 | 5 | 1 | 1 | 0 |
| Nicolas *et al.* 'sliterature review (2) | 0 | 3 | 4 | 1 | 0 | 1 |
| RefSeq Non-Coding (2) | 63 | 144 | 3 | 1 | 0 | 1 |
| Rfam, conservative (2) | 107 | 90 | 10 | 4 | 3 | 0 |
| BsubCyc Non-Coding (3) | 38 | 136 | 6 | 2 | 1 | 1 |
| Dar *et al.* riboswitches (3) | 25 | 9 | 15 | 27 | 6 | 0 |
| Nicolas *et al.* predictions (4) | 111 | 0 | 3 | 1 | 0 | 0 |
| Rfam, medium (4) | 112 | 94 | 13 | 4 | 3 | 1 |

**Table S3.** Distances to the nearest annotation in the merged gene set compared to the Nicolas *et al.* predicted UTRs and intergenic regions. Because Nicolas *et al.* separate UTRs into non-overlapping elements, we added the lengths of the fragments to better convey the length of the biological region.

| distance to closest gene | 3' UTR | 5' UTR | intergenic | internal |
| --- | --- | --- | --- | --- |
| Overlapping | 36 (14%) | 93 (14%) | 23 (7%) | 13 (7%) |
| 0..100 | 38 (15%) | 210 (31%) | 77 (24%) | 81 (44%) |
| 100..500 | 78 (31%) | 320 (47%) | 163 (51%) | 83 (45%) |
| 500..1,000 | 53 (21%) | 39 (6%) | 34 (11%) | 8 (4%) |
| 1,000..2,000 | 37 (15%) | 7 (1%) | 18 (6%) | 1 (1%) |
| 2,000+ | 7 (3%) | 7 (1%) | 4 (1%) | 0 |

**Table S4.** Overlap based comparison of our computed UTRs without those annotated by Nicolas *et al.* including their intergenic regions. We consider overlaps of at least 25 bp. We indicate the number of BSGatlas UTRs that do not overlap tiling-array UTRs and are thus new in the last column. The BSGatlas does not annotate intergenic regions, yet those annotated in Nicolas *et al.* do overlap with BSGatlas UTRs, which are shown. We indicate for each overlap if the types differ.

| Description | Nicolas et al | BSGatlas | new UTRs |
| --- | --- | --- | --- |
| **5'UTR** |  |  |  |
| Annotated | 676 | 2,825 | 2,057 |
| Overlap with same type | 632 |  |  |
| Without/Low overlap | 39 |  |  |
| Overlaps internal_UTR | 5 |  |  |
| **3'UTR** |  |  |  |
| Annotated | 249 | 1,642 | 1,315 |
| Overlap with same type | 231 |  |  |
| Without/Low overlap | 17 |  |  |
| Overlaps 5'UTR | 1 |  |  |
| **internal UTR** |  |  |  |
| Annotated | 186 | 1,126 | 898 |
| Overlap with same type | 138 |  |  |
| Without/Low overlap | 35 |  |  |
| Overlaps 5'UTR | 6 |  |  |
| Overlaps 3'UTR;5'UTR | 4 |  |  |
| Overlaps 3'UTR | 3 |  |  |
| **intergenic** |  |  |  |
| Annotated | 319 | N/A | N/A |
| Without/Low overlap | 166 |  |  |
| Overlaps internal_UTR | 85 |  |  |
| Overlaps 3'UTR | 49 |  |  |
| Overlaps 5'UTR | 13 |  |  |
| Overlaps 3'UTR;5'UTR | 6 |  |  |

**Figure S1.**
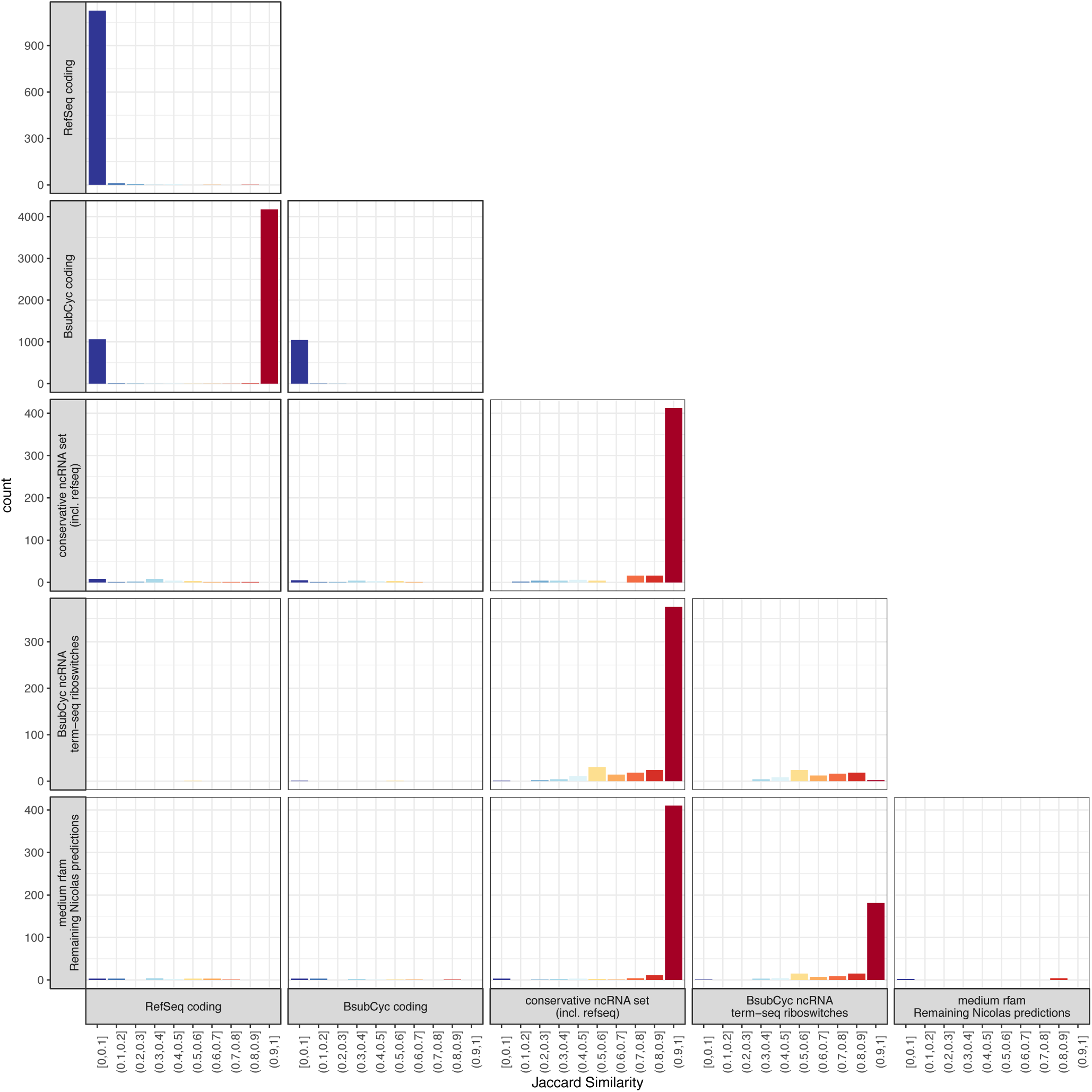
Distribution of Jaccard similarities between all overlapping pairs of genes from the collective annotation, separated by resource. Similarities between each overlapping gene pair (coding and non-coding), identity is ignored. The staining of the histogram bars were added as visual aids.

**Figure S2.**
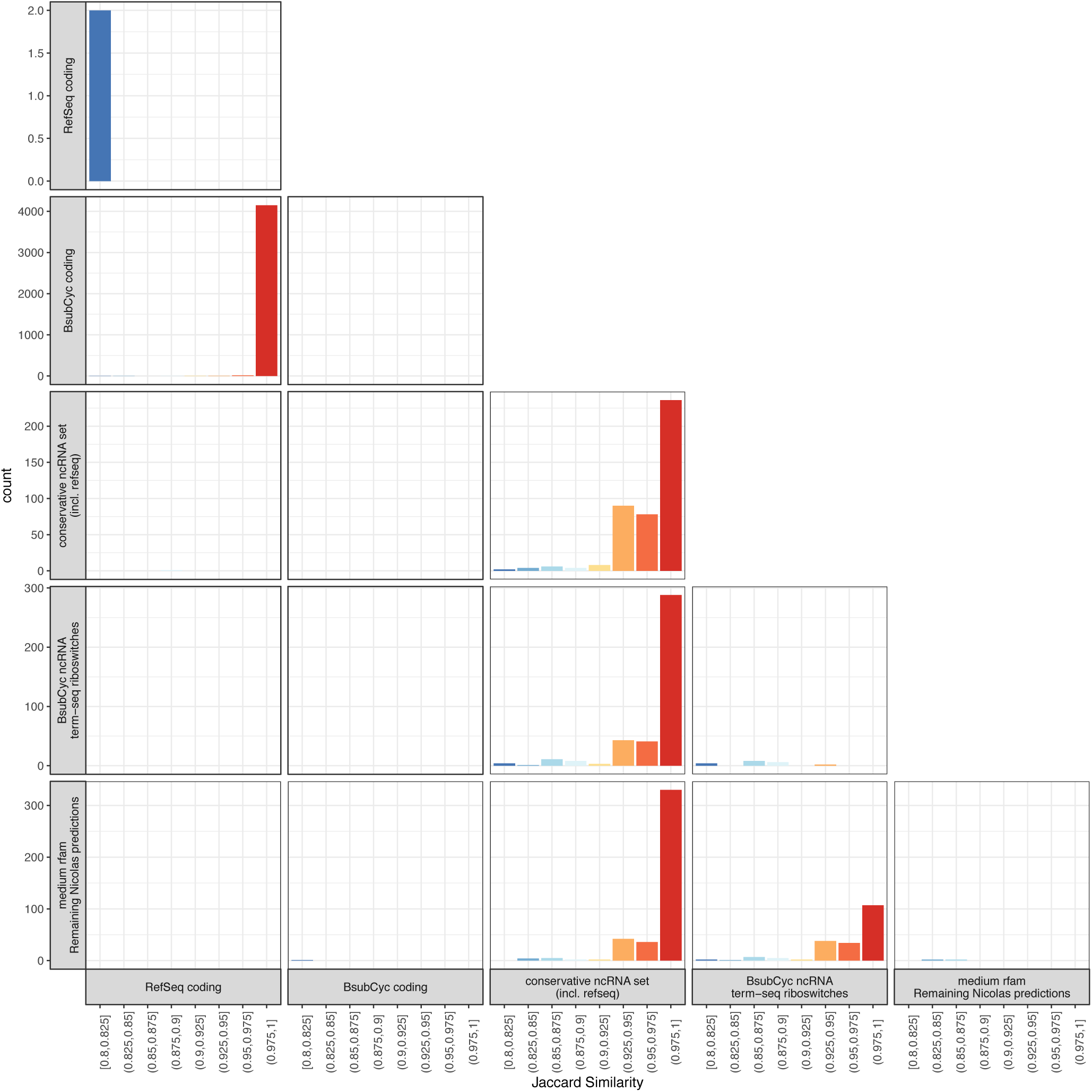
Distribution of Jaccard similarities between all overlapping pairs of genes from the collective annotation, separated by resource. Shown are only similarities above 80%. Similarities between each overlapping gene pair (coding and non-coding), identity is ignored. The staining of the histogram bars were added as visual aids.

**Figure S3.**
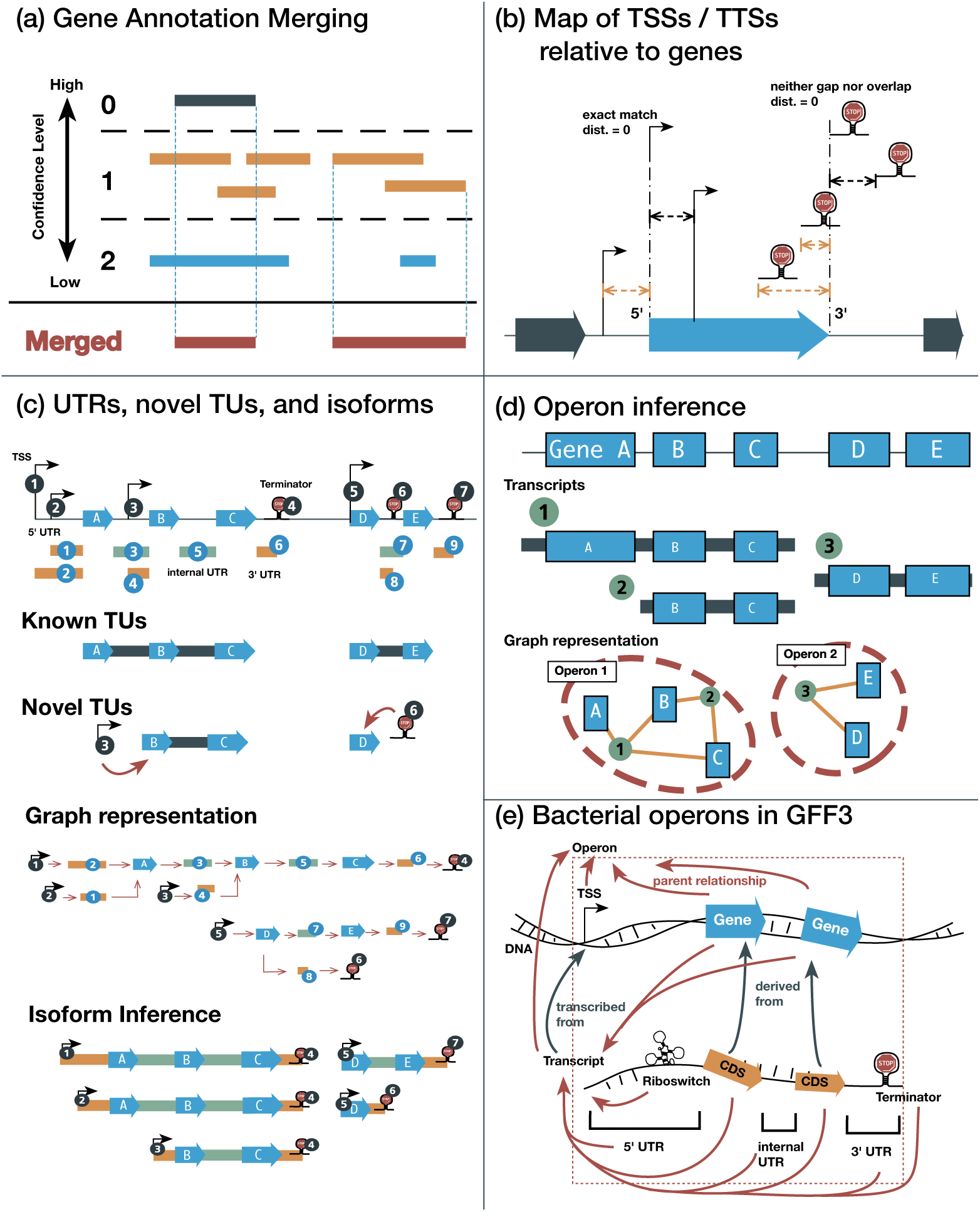
Outline of the annotation creation procedure. (a) Gene annotation merging. Shown are two genes for which the annotation resources provide differing coordinates. The merged coordinates are taken from the highest confidence level, or the union if there are multiple. (b) Distances that are used to determine the transcription start sites (TSSs) and terminator sites (TTSs) map. The TSS distances are relative to the 5’ end of a gene, for a TTS to the 3’. Instead of a single nucleotide position, TTSs annotated an interval, such that the distances are computed as shown. The orange highlighted distances are notated as a negative value. (c) Computation of untranslated regions (UTRs), novel transcriptional units (TUs), and transcripts. Given a TSS/TTS map, 5’ and 3’ UTRs were placed in the space between them and the associated up-/down-stream gene. Internal UTRs were implied by known TUs. Novel TUs are implied by a TSS or TTS that is associated with a gene, which is either not the first or last gene in direction of transcription. The full isoform list is inferred from all paths between TSSs and TTSs, which we derived from a graph. (d) Operon inference. We derived operons by finding connected components in a graph with the transcripts and genes as nodes and edges indicating which genes are transcribed by which transcript. (e) Bacterial operons in GFF3. The GFF3 format models bacterial operons as shown: Each operon/UTR/gene/structure is an entry in the file, although each gene also has an extra entry to represent the transcribed region. The relationships between the entries are noted as indicated by the arrows.

**Figure S4.**
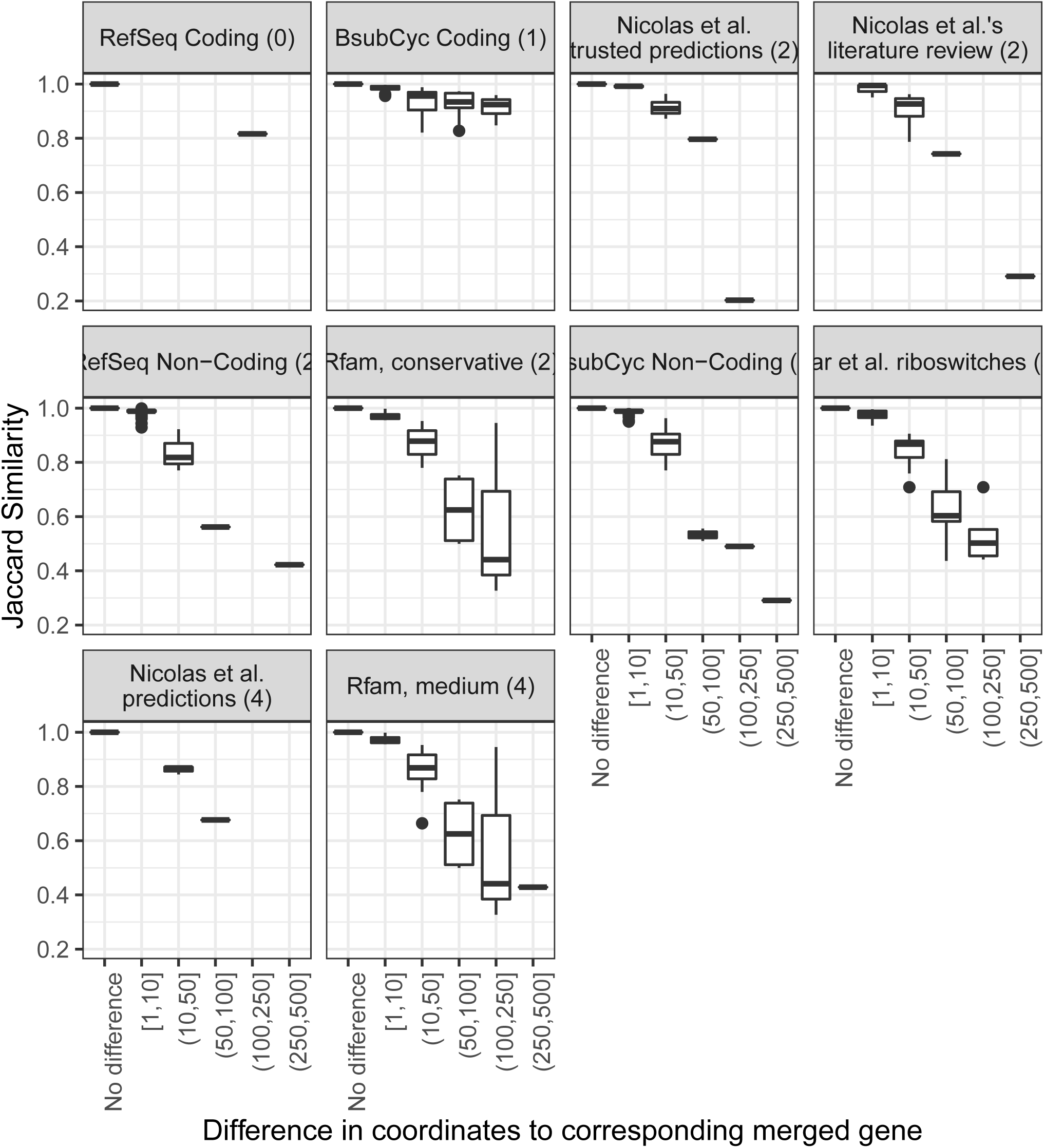
Comparison of the coordinates from each gene annotation resource with those from resulting genes after merging. Shown are the distributions of Jaccard similarity for various ranges of absolute coordinate differences in nucleotides. The numbers of how often a refinement in absolute numbers occurred are stated in Table S2.

**Figure S5.**
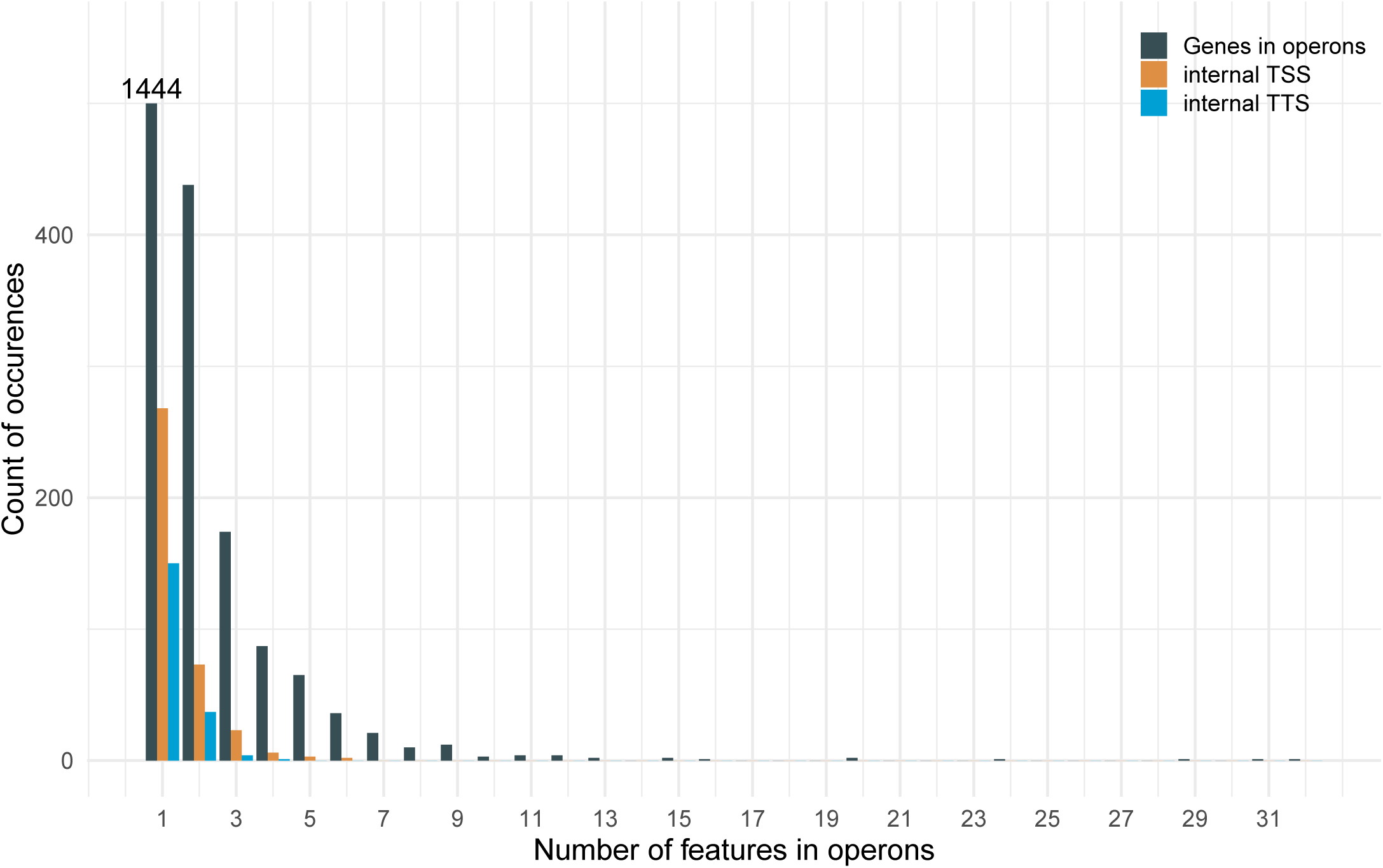
Distributions of the various features, such as the number of genes and internal TSSs / TTSs, for our computed operons in *B. subtilis*.

**Figure S6.**
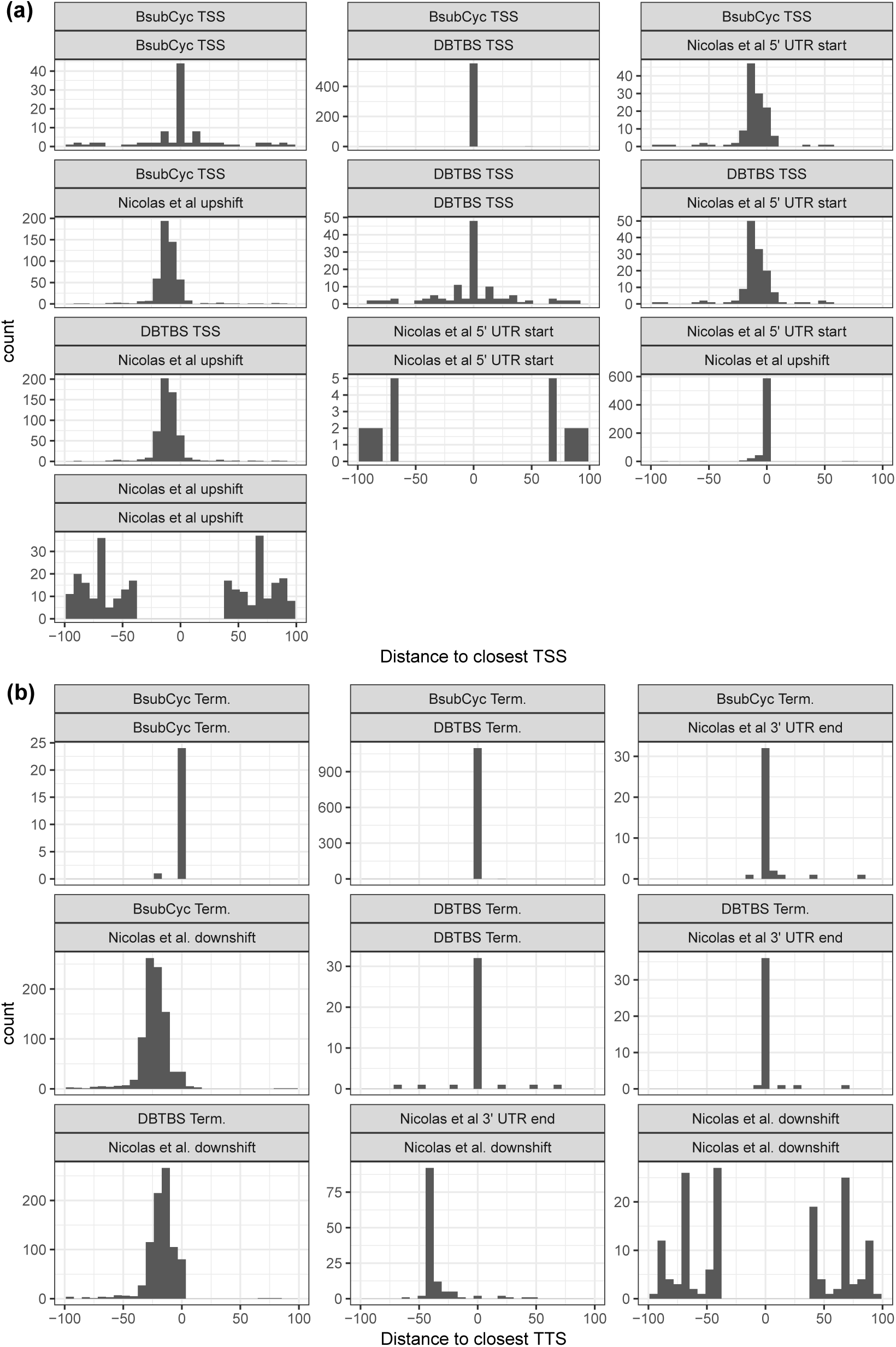
Shown are the histograms of nearest distances between annotated (a) transcription start (TSS) and (B) termination sites (TTS) for the BsubCyc, DBTBS, and Nicolas *et al.* resources. The distances are zero if the corresponding nearest neighbor overlap. A negative distance, such as in the comparison of a TSS from DBTBS versus a Nicolas et al upshift, indicates that the DBTBS annotated TSS is in direction of transcription after/downstream of the upshift. Only absolute distances below 100 bp are shown.

**Figure S7.**
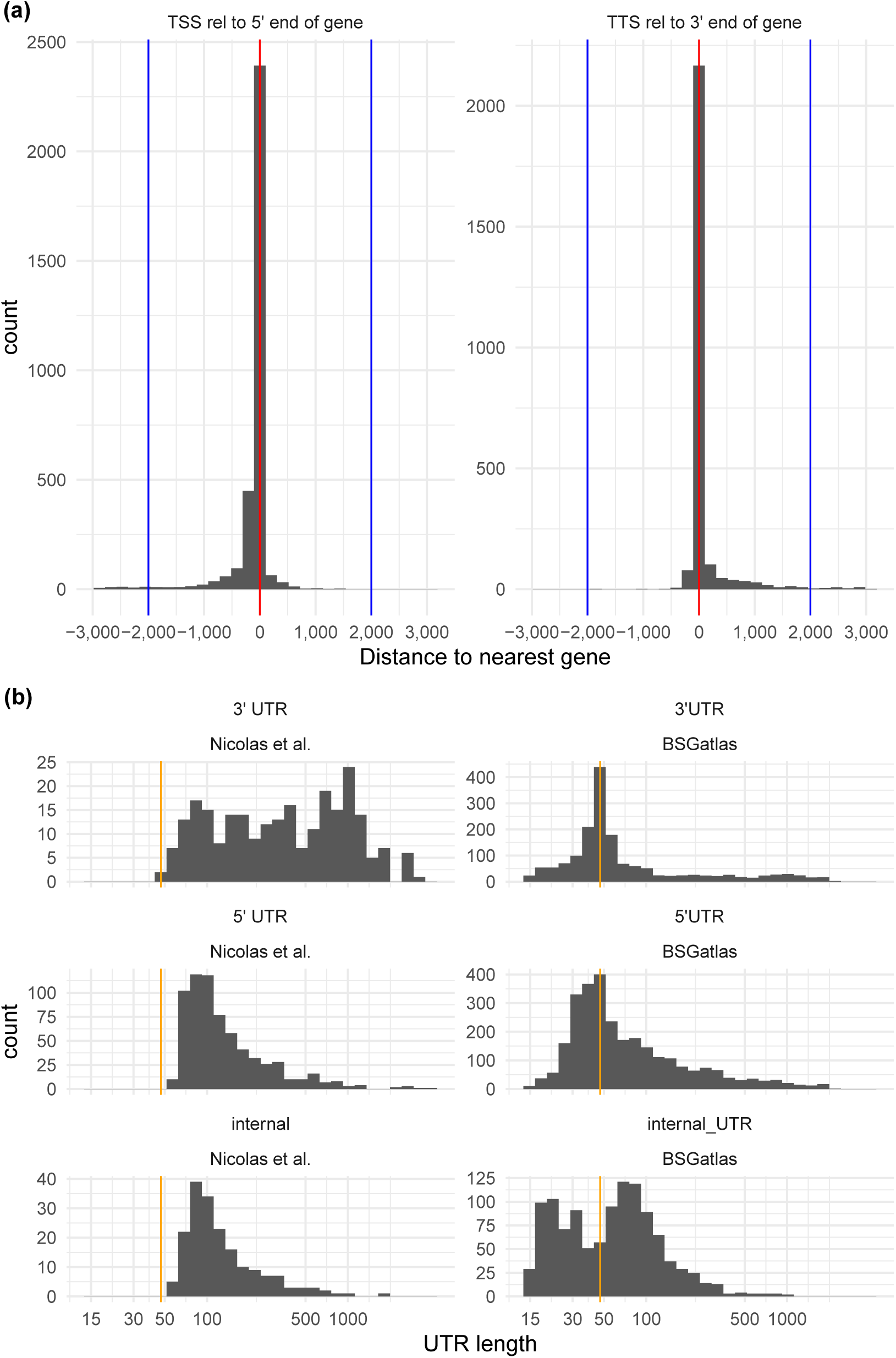
(a) Distribution of TSSs and TTSs relative to the closest 5’/3’ end of genes indicated in red. The blue lines indicate the cut-off we chose for the computations of UTRs. These distances are computed as shown in Figure S3b. (b) Distribution of lengths of our computed UTRs in comparison to those found in Nicolas *et al.*’s tiling-array study. The UTRs of the latter resources have a minimal length of 47, which is indicated with the orange line.

**Figure S8.**
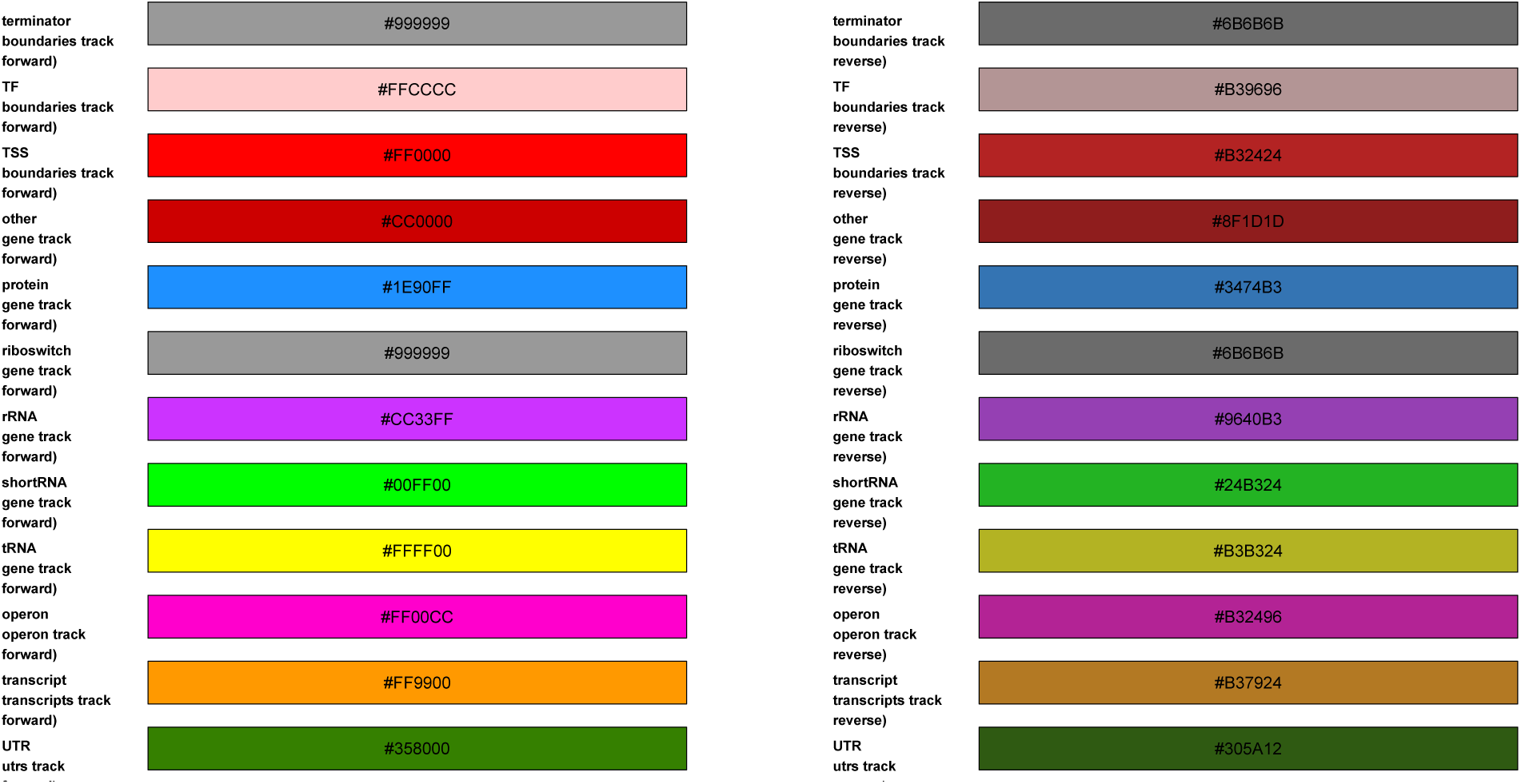
The color scheme for each type of the different annotated element (genes, structures, binding sites). Elements that are on located on the reverse strand are shown in a slightly darker color. We use this color coding across the different annotation visualizations that we offer in the UCSC browser hub, the GFF3 file, and the quick browser on the gene detail pages. Similar looking pairs of color or possibly for color blindness disadvantageous were avoided by putting these on separate gene tracks.

